# An alpha- to gamma-motoneurone collateral can mitigate velocity-dependent stretch reflexes during voluntary movement: A computational study

**DOI:** 10.1101/2023.12.08.570843

**Authors:** Grace Niyo, Lama I Almofeez, Andrew Erwin, Francisco J Valero-Cuevas

## Abstract

The primary motor cortex does not uniquely or directly produce alpha motoneurone (*α*-MN) drive to muscles during voluntary movement. Rather, *α*-MN drive emerges from the synthesis and competition among excitatory and inhibitory inputs from multiple descending tracts, spinal interneurons, sensory inputs, and proprioceptive afferents. One such fundamental input is velocity-dependent stretch reflexes in lengthening muscles, which should be inhibited to enable voluntary movement. It remains an open question, however, the extent to which unmodulated stretch reflexes disrupt voluntary movement, and whether and how they are inhibited in limbs with numerous multi-articular muscles. We used a computational model of a *Rhesus Macaque* arm to simulate movements with feedforward *α*-MN commands only, and with added velocity-dependent stretch reflex feedback. We found that velocity-dependent stretch reflex caused movement-specific, typically large and variable disruptions to arm movements. These disruptions were greatly reduced when modulating velocity-dependent stretch reflex feedback (i) as per the commonly proposed (but yet to be clarified) idealized alpha-gamma (*α*-*γ*) co-activation or (ii) an alternative *α*-MN collateral projection to homonymous *γ*-MNs. We conclude that such *α*-MN collaterals are a physiologically tenable, but previously unrecognized, propriospinal circuit in the mammalian fusimotor system. These collaterals could still collaborate with *α*-*γ* co-activation, and the few skeletofusimotor fibers (*β*-MNs) in mammals, to create a flexible fusimotor ecosystem to enable voluntary movement. By locally and automatically regulating the highly nonlinear neuro-musculo-skeletal mechanics of the limb, these collaterals could be a critical low-level enabler of learning, adaptation, and performance via higher-level brainstem, cerebellar and cortical mechanisms.

**Significance:** Muscles have velocity sensors controlled by *γ*-MNs that produce stretch reflexes which could disrupt voluntary limb movements. Whether and how severely those unmodulated stretch reflexes disrupt voluntary movement remains unclear, especially in realistic multi-articular limbs. Our neuromechanical simulations demonstrate that unmodulated stretch reflexes greatly disrupt movements. Modulating the stretch reflex by implementing an idealized version of a long-posited (but yet unclear) *α*-*γ* co-activation greatly mitigates those perturbations. However, a collateral from the *α*-MN to the *γ*-MN (which has been reported among motoneurones but not interpreted in this way) achieves similar functionality. Our results suggest this modulation of the intensity of the stretch reflex by the *α*-MN collateral provides an effective mechanism to locally stabilize the disruptions from stretch reflexes.

## Introduction

The ‘fusimotor system’ provides proprioceptive feedback signals that are important for kinesthesia, posture, balance [1–3], muscle tone [4], and control of voluntary movement [2, 5]. In mammals, it consists of the ‘muscle spindle’ mechanoreceptors and their associated *secondary* (II) and *primary* (Ia) sensory neurons, which sense muscle fiber length and velocity. The intrafusal muscle fibers of the muscle spindle are innervated by the specialized *γ*_*static*_ and *γ*_*dynamic*_ motoneurones to regulate their sensitivity to muscle (extrafusal fibers) length and velocity, respectively [6–9]. It is often suggested that dysregulation of the fusimotor system is responsible for movement disorders such as hyperreflexia, spasticity, dystonia, etc. [10, 11]. However, the regulation and contribution of this fusimotor system to voluntary movements and movement pathologies remain debatable [11].

The fusimotor system first appeared in amphibians and reptiles, but as a simpler ‘skeletofusimotor system’. In that primitive system, *β*-MNs innervate both the bulk of the extrafusal fibers *and* the spindle’s intrafusal fibers, with no *γ*-MNs present [12, 13]. Detailed simulations of the frog hind limb have demonstrated that in these amphibian *β*-MN arrangements adjustment of gain and phase based on muscle spindle inputs suffice to produce accurate swiping movements [14].

The question remains, however, what the evolutionary pressures could have been to drive the development of separate and independent *α* and *γ* motoneurones in the mammalian fusimotor system (which has precious few *β*-MNs [9, 13]). A strong case has been made that this ‘complication’ allows the flexible modulation of limb impedance [15–17]. This complication, however, comes at the price that the velocity-dependent Ia signal—if not properly modulated—can be considered a form of ‘internal perturbation’ where stretch reflexes in lengthening muscles (i.e., eccentrically contracting or ‘antagonist’ in the single-joint system) can disrupt or stop joint rotations induced by the shortening muscles (i.e., concentrically contracting or ‘agonists’ in the single-joint system) [4, 8, 18, 19]. When exploring this issue, Granit reported that the activity in *α* and *γ* motoneurones is tightly coupled in a given motoneurone pool, and termed it ‘alpha-gamma linkage’ [20] (now known as ‘*α*-*γ* co-activation’). But the details of how *α*-*γ* co-activation is implemented and regulated remain unclear [21, 22], and are difficult to study due to the experimental challenges of recording from small *γ*_*dynamic*_ and *γ*_*static*_ motoneurones in behaving animals and humans [23–28]. Moreover, the popular but simplified agonist-antagonist conceptual framework is difficult to generalize to limbs driven by numerous multi-articular muscles where the roles of agonist and antagonist become unclear and can change during the movement [18, 19, 29, 30].

In this study, we apply first principles to address two issues. First, in what ways does positive homonymous muscle velocity feedback (i.e.,velocity-dependent stretch reflexes) perturb 3-dimensional arm movements in the general case of numerous multi-articular muscles? And second, how does modulation of velocity-dependent stretch reflex gains in the mammalian fusimotor system mitigate these disruptions? We find that unmodulated, physiologically tenable monosynaptic velocity-dependent stretch reflexes do, in fact, disrupt voluntary movements in significant, variable and task-specific ways. Moreover, both idealized *α*-*γ* co-activation and a simpler *α*-MN collateral projection to *γ*-MNs can greatly reduce disruptions for most voluntary movements. We propose that such previously unrecognized collaterals could be a low-level enabler of learning, adaptation, and performance that can be evolutionarily and developmentally complemented by *α*-*γ* co-activation and other brainstem, cerebellar and cortical mechanisms.

## Results

### Unmodulated velocity-dependent stretch reflexes cause large, variable disruptions of the endpoint trajectory in task-dependent ways

We simulated 1,100 different *α*-MN coordination patterns to study how unmodulated velocity-dependent stretch reflex disrupt movement trajectories in a 25-muscle, 5-degree-of-freedom arm model of a *Rhesus Macaque* monkey—and how the disruptions change with different spindle sensitivity levels (i.e., increasing velocity-dependent stretch reflex gain, *k*). The neural circuit, schematic diagram, sample *α*-drive (coordination patterns or activation signals) to muscles, and resulting open-loop reference trajectory are shown in the top row of Fig. 1. The same are shown for the closed-loop simulation with unmodulated reflexes (at ten different gains) in the bottom row. Each of the 25 afferented muscles consists of a simple muscle spindle model that outputs positive velocity of lengthening muscles (i.e., velocity of stretch) as afferent feedback to its *α*-MN, subject to the reflex gain *k*.

Our 1,100 open-loop simulations of arm endpoint trajectories resulted in small and large arm movements (*SI Appendix*, Fig. S1), which were disrupted when closing the loop with the velocity-dependent stretch reflex. Unmodulated reflexes resulted in disrupted *movement trajectories* (e.g., Fig. 2, cases 635, 147, 430, 884, and 122). Conversely, in other arm movements, the terminal positions remained unaffected by the velocity-dependent stretch reflex (e.g., Fig. 2, cases 5,518 and 596). Additionally, increase in reflex gain could change the movement direction (e.g., Fig. 2), case 884 and 122). In all arm movements, the disruptive effect was consistently visible even at gain *k* = 1. (Figs. 2, 4A, and 5A) and increased when the reflex gain was increased; however, the nature of the disruption in the trajectory and endpoint differed across movements. The results show that the disruption in the arm endpoint trajectory depended on both the stretch reflex gain and the movement itself. This is because even similar movements can induce different muscle fiber velocity profiles [19].

**Fig 1.**
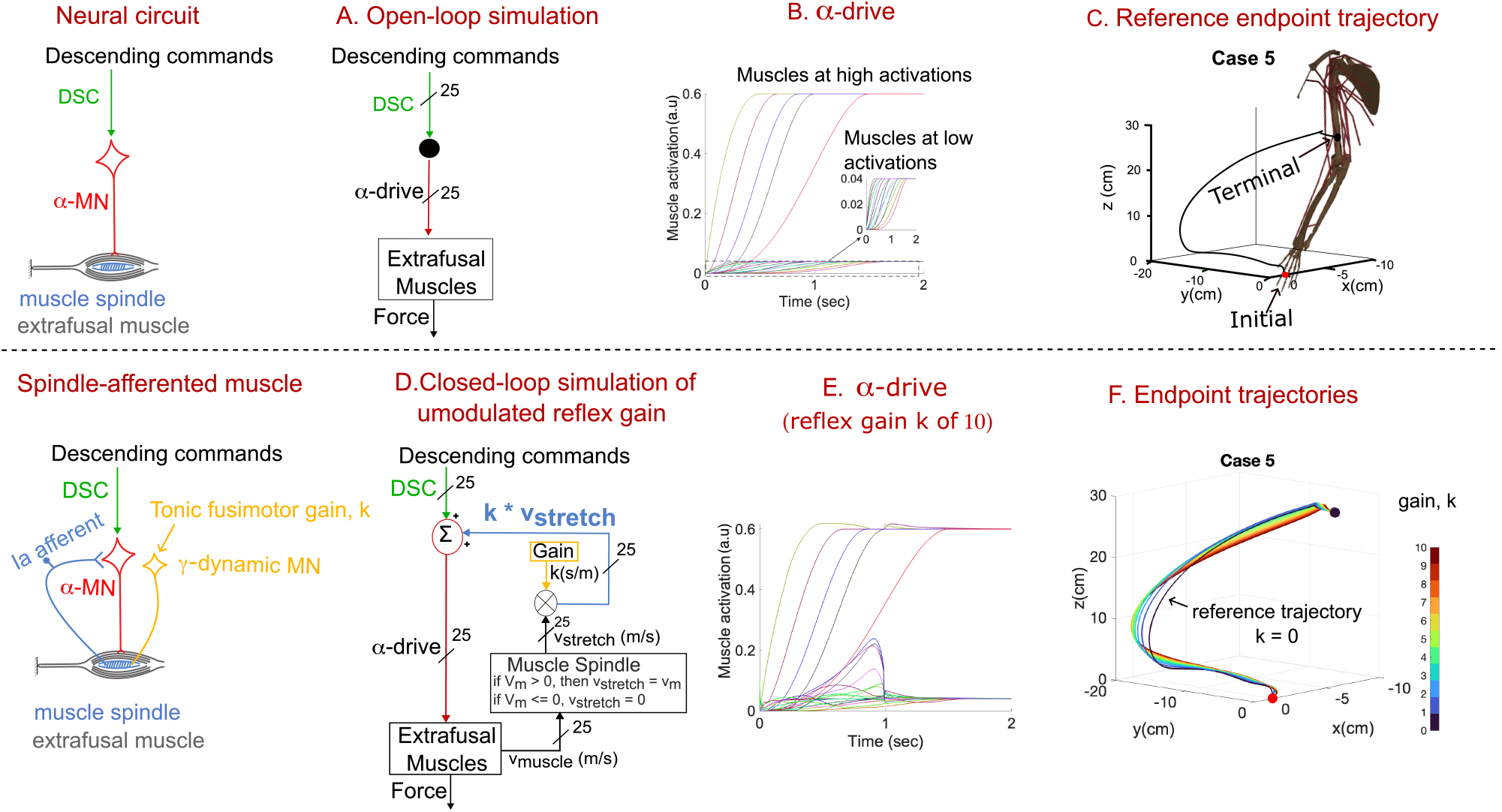
Neural circuit, simulated circuit, [*α*-drive to muscles, and resulting endpoint trajectories for case 5 out of 1,100. **Top row:** For the open-loop reference trajectory. **Bottom row:** For the unmodulated velocity-dependent stretch reflex simulations at 10 gain levels, *k*. **(A)** Open-loop simulation of arm movement without velocity-dependent stretch reflex for the neural circuit shown. **(B)** Sample of *α*-drive to muscles, and **(C)** the ensuing reflex-free reference trajectory of the endpoint (distal head of the third metacarpal) from the initial position (red dot) to the terminal position (black dot). When closing the velocity-dependent stretch reflex loop, **(D)**, the muscle stretch velocity was multiplied by a reflex gain *k* to produce the unmodulated velocity-dependent stretch reflex feedback (*k*\**V*_*stretch*_). Sample *α*-drive to muscles at a reflex gain *k* = 10 is shown in **(E)**, with the resulting endpoint trajectories in **(F)** color-coded to reflex gains from *k* = 0 (i.e., open-loop reference endpoint trajectory) to *k* = 10. See details in the Methods section.

**Fig 2.**
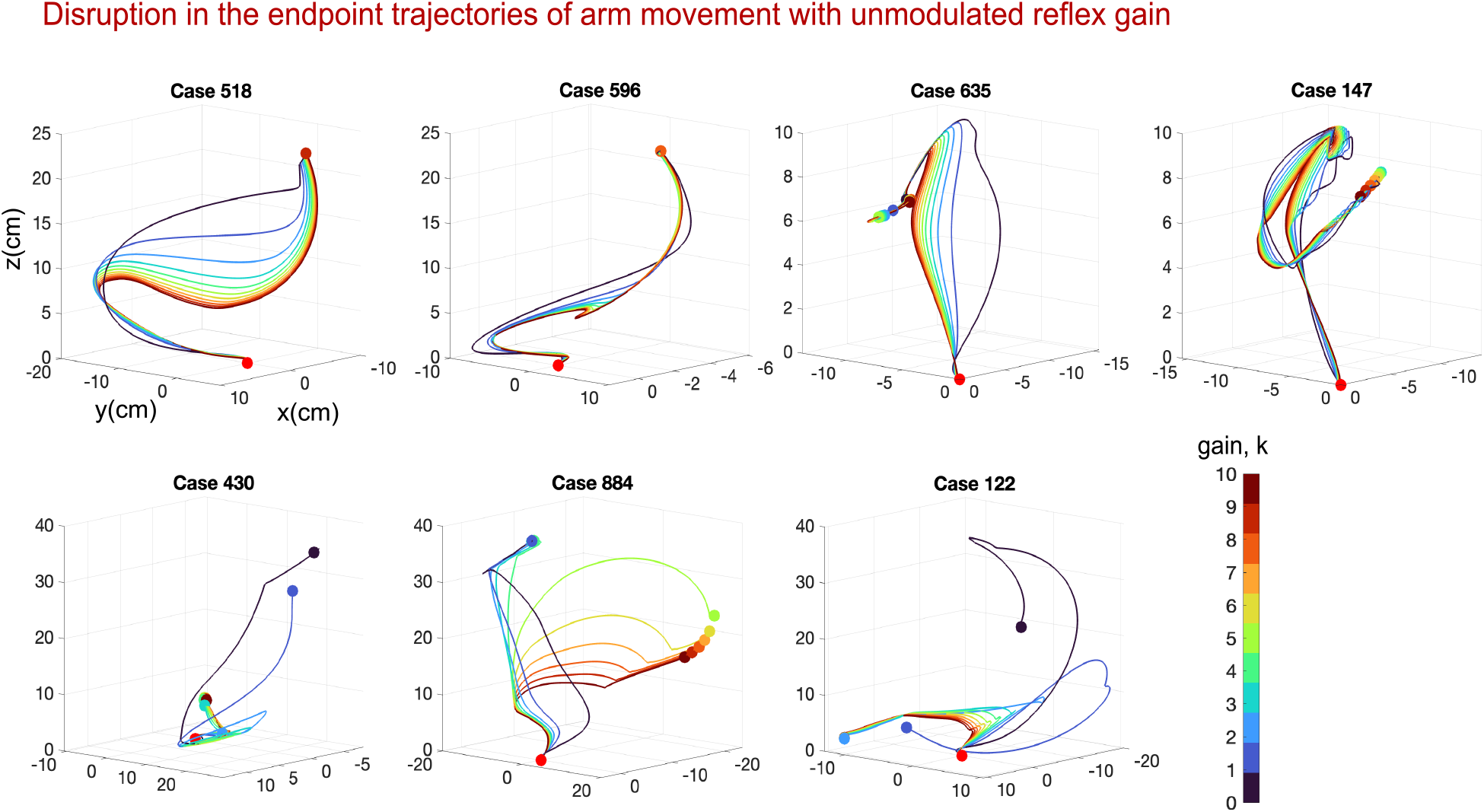
Unmodulated reflex gains can cause large, variable, and movement-specific disruptions. Seven examples, in addition to an eighth shown in Fig. 1F, show representative disruptions of the endpoint trajectory caused by unmodulated velocity-dependent stretch reflex. Increasing the reflex gain *k* progressively disrupts the endpoint trajectory in different ways. In case 5 (Fig. 1F) and cases 518 and 596, only the movement trajectories were disrupted. In cases 635, 147, 430, 884, and 122, both the movement trajectory and terminal endpoint position were disrupted; in cases 884 and 122, increasing the reflex gain *k* even changed the movement direction. Figures 3A and 3B show the same examples when the velocity-dependent stretch reflex feedback was modulated as per idealized *α*-*γ* co-activation and scaled via an homonymous *α*-to-*γ* collateral, respectively.

### Idealized *α*-*γ* co-activation and a collateral from the *α*-MN axon to *γ*-MNs reduce movement disruption caused by velocity-dependent stretch reflex

We then investigated how the disruption in movement trajectory changes when velocity-dependent stretch reflex were modulated as per idealized *α*-*γ* co-activation (Fig. 3A). We also explored an alternative modulation of *γ*-MN activity. It has been known for some time that motoneurones in general have collateral branches (i.e., emanating from their axons) projecting to other motoneurones [31–34, 34]. The most well-known is the collateral to a reciprocally self-inhibiting Renshaw cell [35, 36]. However, we posited such a collateral could also project to homonymous *γ*-MNs. We implemented this homonymous *α*-to-*γ* collateral as a scaling of the velocity-dependent stretch reflex proportional to the *α*-MN output (Fig. 3B). We find the disruptions to the arm trajectories became small at all reflex gains when the simulated stretch reflexes were modulated as per idealized *α*-*γ* co-activation or scaled by the homonymous *α*-drive (Fig. 3). Physiologically, such modulation is in effect *homonymous* inhibition of *γ*-MNs—which is distinct from the well-known *reciprocal* inhibition of the ‘antagonist’ *α*-MNs in the single-joint system [37, 38]. This is achieved numerically in our simulations by a multiplication by a value less than 1 (i.e., the *α*-MN drive) in both idealized *α*-*γ* co-activation and homonymous *α*-to-*γ* collateral. Notwithstanding, Figures 4B and 4C show four examples of endpoint trajectories that retained large disruptions even after reflex modulation. Thus reflex modulation is not a panacea (see Discussion).

**Fig 3.**
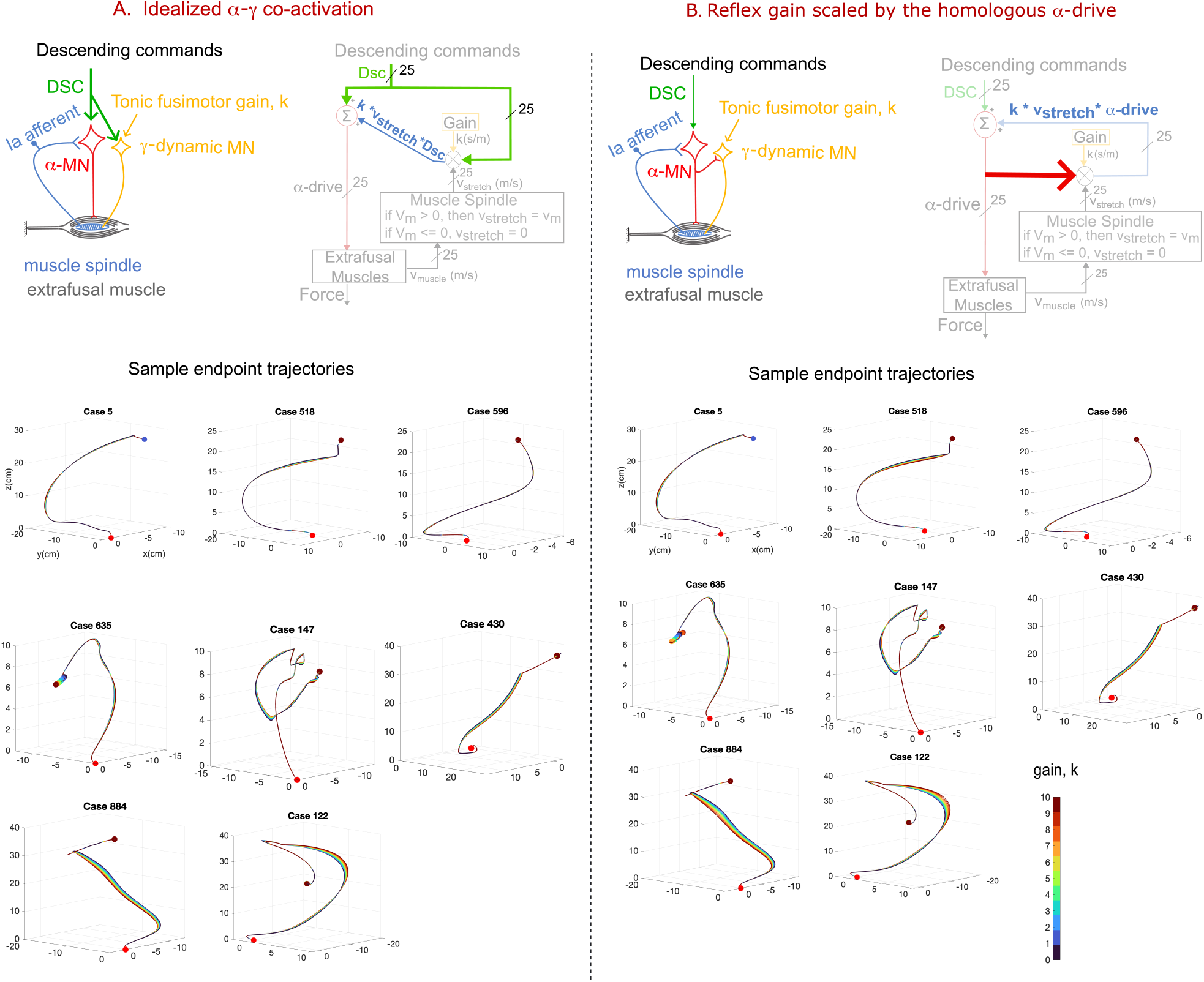
Modulating velocity-dependent stretch reflex feedback reduces disruptions in the arm movement trajectory. **(A)** Stretch reflex modulated as per idealized *α*-*γ* co-activation. The descending *pre-synaptic* command (DSC, green color) to *α*-MN scaled the reflex gain *k*, which resulted in velocity-dependent stretch reflex feedback equal to *k*\**V*_*stretch*_*DSC. **(B)** For the homonymous *α*-to-*γ* collateral (red bold arrow), the reflex gain was scaled by the muscle’s *post-synaptic α*-MN drive to produce velocity-dependent stretch reflex feedback equal to *k*\**V*_*stretch*_\**α*-drive. We show the same seven examples shown for unmodulated reflexes in Figures 1F & 2, see Methods. Analysis of the effects of velocity-dependent stretch reflex and their modulation on the arm movement trajectories of all 1,100 arm movements is shown Figures 5 B & C at each reflex gain. Furthermore, quantitative and statistical comparison of the disruption in the endpoint trajectories for idealized *α*-*γ* co-activation and homonymous *α*-to-*γ* collateral provided in *SI Appendix*, Fig. S2 and Figure 6.

**Fig 4.**
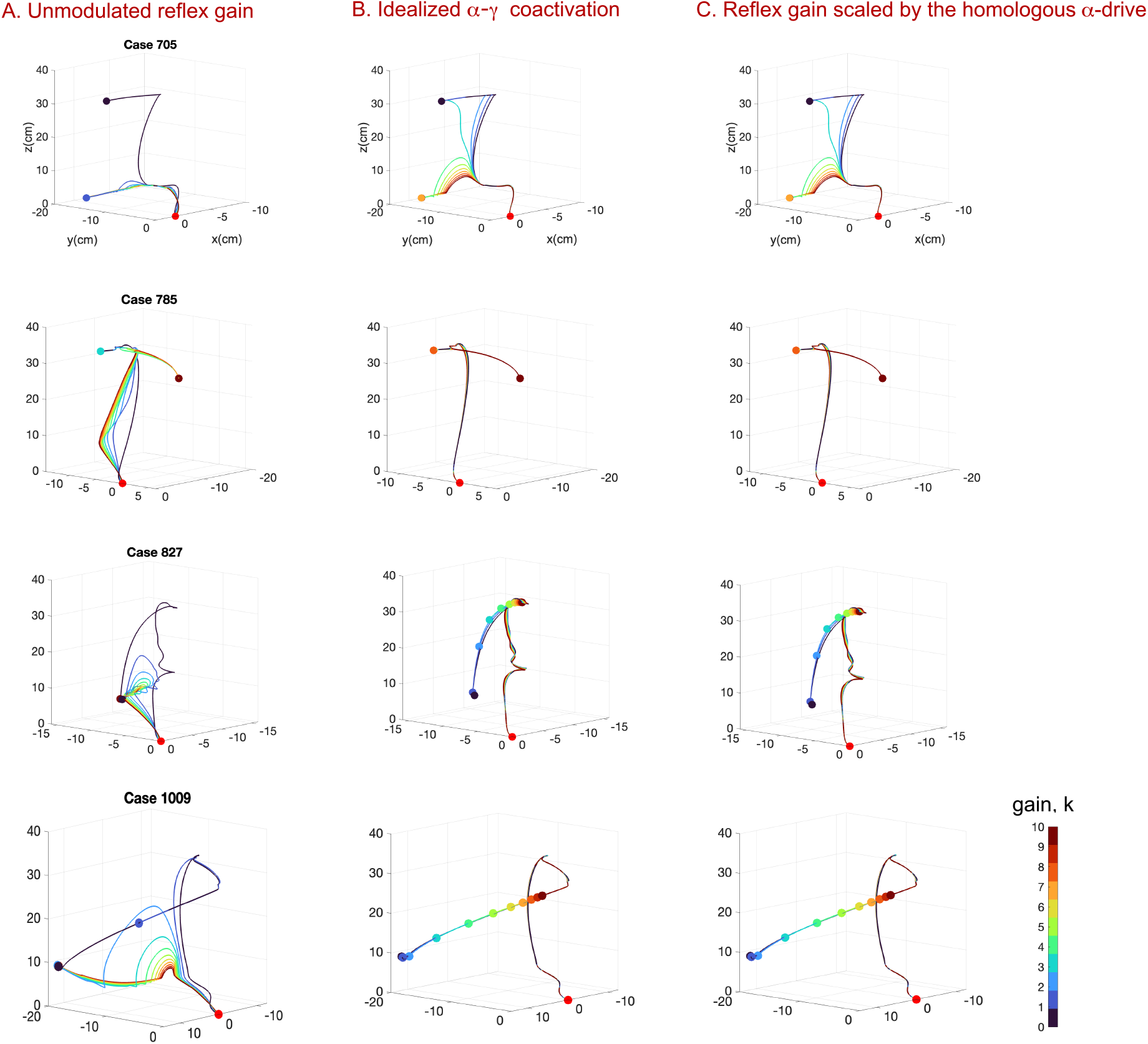
Reflex modulation is not sufficient in all cases. We show four sample cases **(A)** where the movement disruption remained large even after idealized *α*-*γ* co-activation **(B)** or homonymous *α*-to-*γ* collateral **(C)**.

In Figure 5, we show that idealized *α*-*γ* co-activation and homonymous *α*-to-*γ* collateral both generally reduce disruptions across all 1,100 arm moments, where the cumulative residual and terminal error naturally increased at higher reflex gains. Our statistical analysis revealed a significant decrease in movement disruptions with reflex modulation (Fig. 5B or 5C, p *<* 0.001). However, no significant differences were found in the terminal errors between idealized *α*-*γ* co-activation and homonymous *α*-to-*γ* collateral (p *>* 0.057, bottom rows in Fig. 5B & 5C respectively). We did find statistically significant (p *<* 0.001), but numerically small (Figure 6), differences in cumulative residual between them (top rows in Fig. 5B & 5C). The distribution of the cumulative residual and terminal error across all movements at each reflex gain *k* are shown in *SI Appendix*, Fig. S2. Furthermore, Dynamic Time Warping (see Methods) between the endpoint trajectories from idealized *α*-*γ* co-activation and homonymous *α*-to-*γ* Collateral (Figure 6) showed numerically small cumulative differences (*<<* 1 cm)between the trajectories.

**Fig 5.**
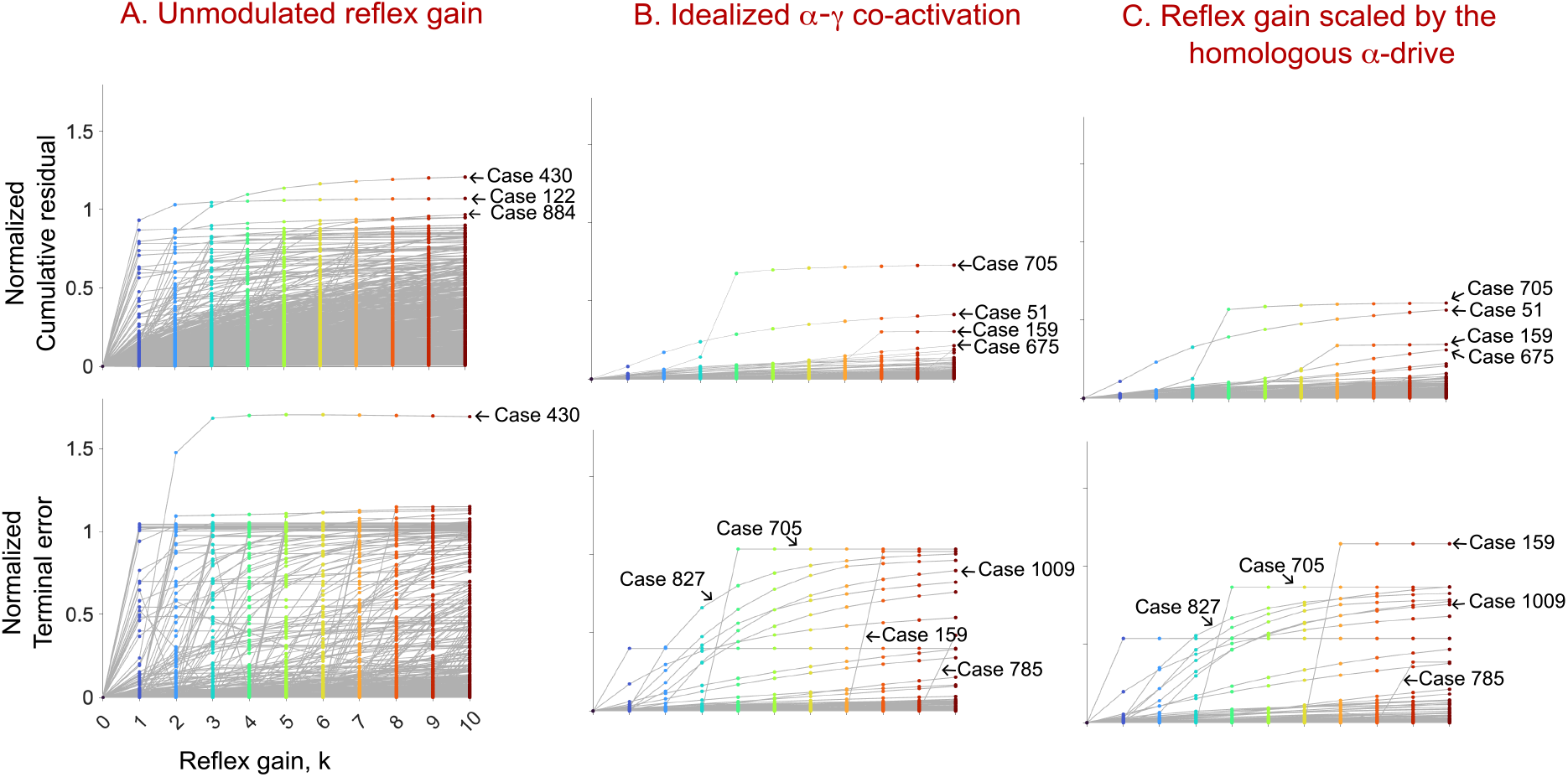
Terminal error and cumulative residual with respect to the reference trajectories were large and variable for the unmodulated stretch reflexes (A); but typically small to negligible when the stretch reflexes were modulated as per idealized *α*-*γ* co-activation (B) or when scaled by the homonymous *α*-drive (C). For each movement, we divided the deviation in movement trajectory (i.e., cumulative residual, CR) and terminal position (i.e., terminal error, TE) by the maximal endpoint displacement of that movement’s reference trajectory(*SI Appendix*, Fig. S1). Both CR and TE (top and bottom plots, respectively) of all 1,100 arm movements at each gain *k* reduced when the reflex gains were modulated.

**Fig 6.**
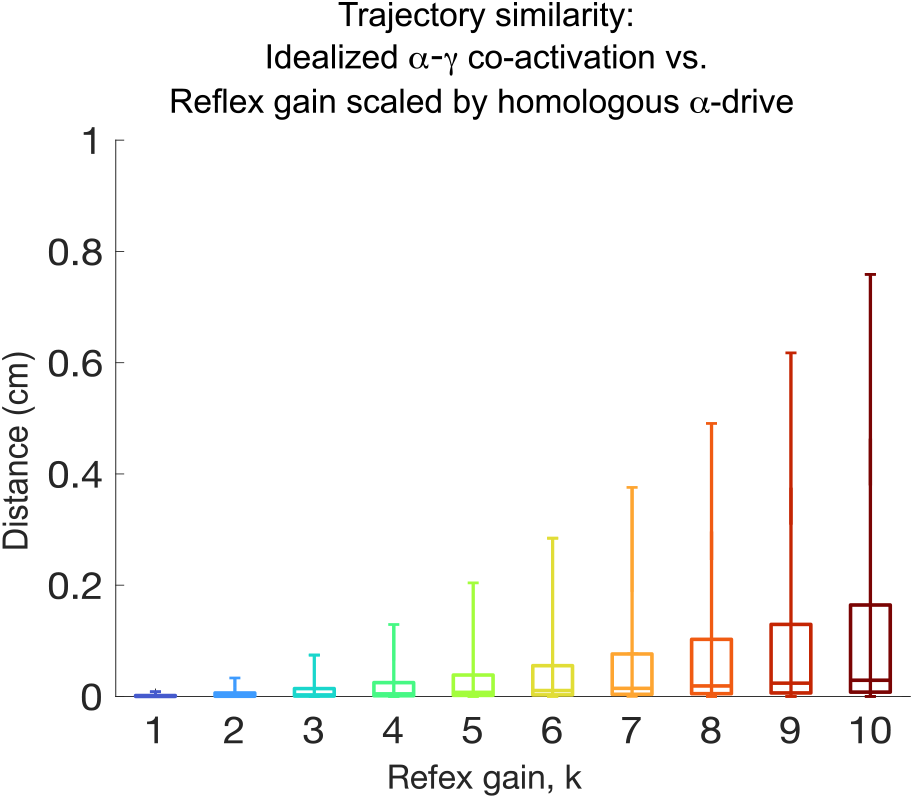
Dynamic Time Warping shows small differences in movement trajectories between idealized *α*-*γ* co-activation and homonymous *α*-to-*γ* collateral. The differences were minimal (*<* 1 cm) even though they increases with gain *k*. For reference, the median endpoint displacement was 21.74 cm, Figure *SI Appendix*, Fig. S1.

For each closed-loop simulation, we computed the peak change in *α*-drive to muscles caused by velocity-dependent stretch reflex feedback (Fig. 7). We found peak changes in *α*-drive to muscles were comparable to those observed in the human arm during interactions with destabilizing environments [39] (i.e., up to 40% MVC). Sample time series of muscle activation signals with velocity-dependent stretch reflex at a gain of ten are provided in *SI Appendix*, Fig. S3. Thus, we believe the disruptions we report are a realistic *computational* prediction of the neuromechanics of limb movement that are not easily obtained *experimentally* —which is one of the most useful applications of computational modeling [40]. For each movement trajectory, there were significant reductions in peak changes in *α*-drive to muscle between unmodulated (Fig. 7A) and modulated velocity-dependent stretch reflexes (Figs. 7B & 7C) for all muscles (p *<* 0.001, across all gains). However, no statistically significant differences were found between idealized *α*-*γ* co-activation (Fig. 7B) and homonymous *α*-to-*γ* collateral (Fig. 7C) in muscles at either low or high activation at p *<* 0.01 levels (required for Bonferroni correction) for each gain.

**Fig 7.**
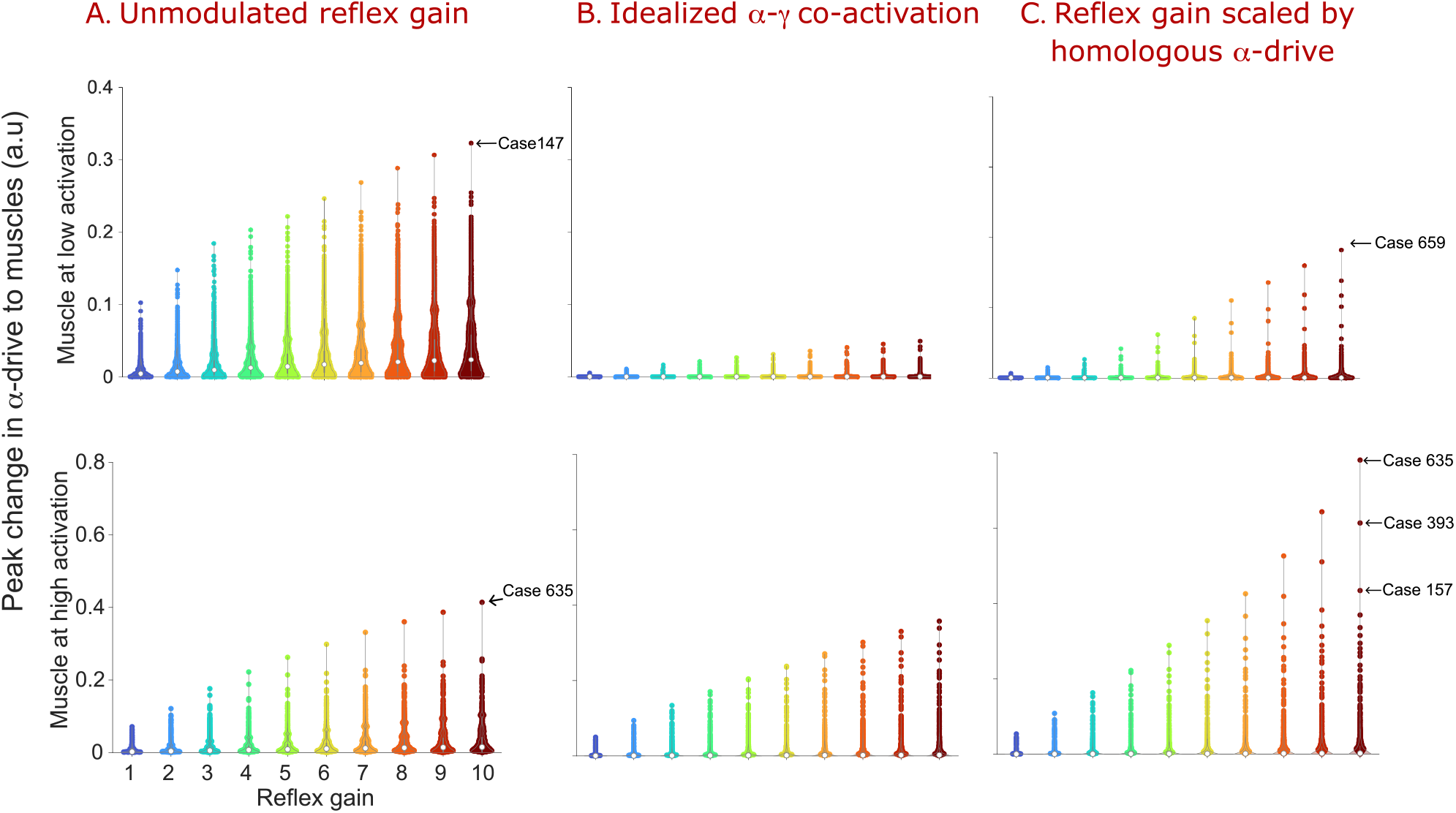
Distribution of the peak change in the muscle activations (normalized to reference feedforward activation) for all 1,100 cases at each reflex gain. The peak change in *α*-drive for unmodulated velocity-dependent stretch reflex **(A)** were significantly larger than those when velocity-dependent stretch reflex were modulated as per idealized *α*-*γ* co-activation **(B)** and homonymous *α*-to-*γ* collateral **(C)**. No statistically significant differences were found between idealized *α*-*γ* co-activation and homonymous *α*-to-*γ* collateral.

## Discussion

We used a computational model of a *Rhesus Macaque* arm with 25 muscles to test whether velocity-dependent stretch reflexes (i.e., simple positive feedback monosynaptic simulating Ia afferents) are sufficiently disruptive to require dynamic modulation to produce accurate movements in realistic multi-articular limbs. We note at the outset that we compare disruptions in 1,100 cases with respect to baseline feedforward movement trajectories produced by randomly-generated muscle activation time histories (Fig. 1). As per our prior work, such computational exploration with randomized trials allows the study of motor control in general—without the need to appeal to, espouse, defend, or justify any form of neural control principle, optimization, or cost function [19, 40–42] We considered these movements to be realistic because the arm’s joints were limited to operate within their natural ranges of motion and the muscle activations produced reach-like movements. Admittedly, not all activations patterns or movements will be seen in a macaque, and non-reaching (i.e., cyclical) movements would necessitate different coordination patterns. However, had we assumed *a priori* a particular control strategy, optimization, set of motor primitives [43], synergistic or Bayesian control law [41, 44], our results could be limited by those assumptions, and would be less general and convincing.

Our general results show that the disruptions of the movements caused by the velocity-dependent stretch reflexes are large, variable, and task-dependent enough to need inhibition, as has been proposed—but never quantified—by Sherrington and others [8, 18, 29, 45, 46]. Frankly, we were surprised by the magnitude and variety of types of disruptions that arose when velocity-dependent stretch reflexes are not modulated. We then demonstrate that both idealized *α*-*γ* co-activation and a plausible homonymous *α*-to-*γ* collateral significantly reduce those disruptions. Note that the modulation of velocity-dependent stretch reflex as implemented in Eqns. 2 & 3 makes ‘*α*-drive’ (a number between zero and 1) an attenuator. Thus they can be thought of as an inhibitor of the velocity-dependent stretch reflex. Importantly, collaterals among MNs have been reported to exist—even by Renshaw himself in addition to the eponymous reciprocally inhibitory neurones [31–33, 47]—but not thought to provide this homonymous modulatory function. We can only speculate about the spinal mechanisms that make reflex intensity track *α*-drive. Figure 8B and 8C propose two schematic circuits that in principle can achieve this, but future work is needed to uncover the actual interactions via excitatory, inhibitory or dis-inhibitory synaptic connections (or the strength of synaptic projections and collateral density) needed to regulate *γ*-drive proportional to the homonymous *α*-drive.

**Fig 8.**
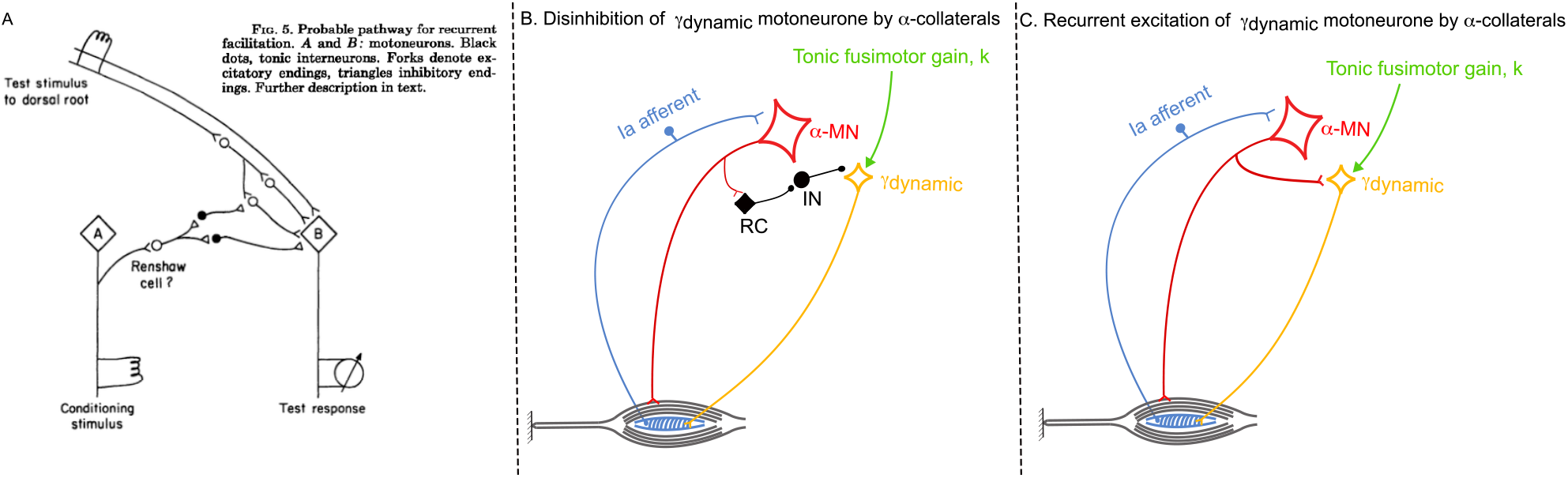
Three schematic homonymous collateral circuits compatible with Eq.3: From Wilson and Burgess (1962) [33] (**A**), its adaptation to have the collateral disinhibit the homonymous *γ*-MN via a disynaptic projection via a Renshaw cell (**B**), and a straightforward monosynaptic modulatory drive to the *γ*-MN compatible with [47] (**C**). All of these versions of collaterals are neuroanatomically and functionally distinct from, and not equivalent to, idealized *α*-*γ* co-activation.

### Muscle afferentation compels us to revisit the foundations of voluntary movement

The maxim apocryphally attributed to Sherrington that *‘inhibition is as important as excitation’* is regularly emphasized in the iconic single-joint system with a single agonist-antagonist muscle pair that customarily introduces students to the spinal motor system [48, 49]. This simplified neuromechanical system clearly shows that, for voluntary joint rotation to occur, the shortening of the ‘agonist’ muscle is made possible by the inhibition of length- and velocity-dependent stretch reflexes of the lengthening ‘antagonist’ muscle. As has been extensively documented in highly controlled experimental single-joint preparations, this can be made possible by propriospinal reciprocal inhibition (within or across limbs), broad branching of proprioceptive signals via inteneuronal pathways, or coordinated descending inhibitory signals [4, 37, 38, 45].

However, how reciprocal inhibition and its circuitry generalize for voluntary movement of realistic *multi-joint limbs with numerous multi-articular muscles* remains an open question in theories and experiments of motor control. As a result, the mechanisms and nature of the modulation of velocity-dependent stretch reflex is left to specialists to grapple with [37, 38]. The reasons are multiple. For example, a same muscle can switch between eccentric and concentric contraction during a same movement, and the roles of agonist and antagonist lose their meaning [18, 19, 50]. More fundamentally, the addition of muscle afferentation to the problem of motor control transforms muscle coordination into a mathematically *over-determined* problem (i.e., there is at most one solution: any single eccentrically contracting muscle that fails to regulate its velocity-dependent stretch reflex can lock or disrupt the movement) [18, 19]. This is the opposite of the traditional view that muscle coordination is mathematically redundant (i.e., *under-determined* where infinite combinations of muscle forces can produce the same joint torques). This dichotomy or apparent paradox arises because limbs are controlled by afferented musculotendons (linear actuators) that can shorten and lengthen. This makes the control of *joint rotations* (i.e., limb motion) mechanically and neurophysiologically distinct from the control of *net joint torques* (i.e., limb forces) [18, 51, 52].

Muscle afferentation is seldom mentioned in canonical reviews of computational theories of motor control, or is assumed to be regulated by predictive or feedback gain scheduling mechanisms, efference copy and internal models [53–55]. Our results provide fruitful research directions by objectively quantifying the consequences of not modulating muscle afferentation, and emphasizing the possibility of an ecosystem for the modulation of velocity-dependent stretch reflexes that ranges from low-level spinal circuits to hierarchical and distributed strategies.

### Idealized *α*-*γ* co-activation effectively mitigates disruptions from velocity-dependent stretch reflexes

The popular and dominant working hypotheses about the modulation of muscle spindle sensitivity [48, 56] revolve around the coordination between *α*- and *γ*-MN activity in a way that allows (i) muscle proprioception and (ii) appropriate eccentric contractions. There is much evidence that the firing among individual motorneurones in isolated rodent spinal cords (without feedback) exhibits very little variance in their timing [57, 58]. The fact that only one activation pattern is seen per motoneurone pool (without feedback) suggests that both *α*- and *γ*-MNs likely fire in similar patterns in these isolated preparations. But the inability to clearly record from, and distinguish, *α*- and *γ*-MNs has impeded conclusive measurements. This synchronicity could be a consequence of either shared input to both *α*- and *γ*-MNs, or homonymous *α*-to-*γ* connectivity (Fig. 3). It remains unknown how much shared presynaptic input is needed to support such tight firing in the motoneurone pools with feedback in an intact behaving animal.

The traditional idealized version of *α*-*γ* co-activation posits that the *γ*_*static*_ MNs that drive the intrafusal fibers of the secondary (II) spindle afferents (sensitive to muscle length) are activated synchronously with *α*-MNs. This prevents the intrafusal muscle fibers from going slack to maintain secondary spindle sensitivity. However, other than preventing slack in intrafusal fibers, *α*-*γ* co-activation does not explicitly address the modulation of intrafusal primary Ia afferents involved in velocity-dependent stretch reflexes [59]. Other theories are variants of *α*-*γ* co-activation, like *Fusimotor Setpoint* which focuses on Ia stretch-sensitivity during learning [26], but does not address their role in the regulation of arbitrary movements after they have been learned. Two other variants posit that fusimotor drive is played out in time. The first as a *Temporal Template* where the modulation frequency of *γ*_*static*_ MNs in shortening muscles increases to expand the dynamic range of spindles during active movements, and the *γ*_*dynamic*_ MNs prime primary afferents to detect the initiation of muscle lengthening and deviations from intended movement trajectory [60]. The second, *Goal-Directed Preparatory Control*, assumes that the stretch reflex gains are modulated according to the predicted spindle activity [17].

There are some limitations to *α*-*γ* co-activation and its variants. For example, they hinge on the assumption that the system has sufficiently accurate knowledge of the time-varying variables that determine musculotendon lengths and velocities (e.g., the current and future states of all muscles, joint kinematics and external forces). Multiple theories have been proposed to provide such future knowledge for known tasks (which is also needed for learning, error correction, response to perturbations, etc.) including efferent copy, internal models, optimal control, synergy control, and Bayesian estimation [41, 54, 61, 62]. In addition, time delays and uncertainty can conspire to pollute such estimates if they rely on the supra-spinal processing of sensory signals to estimate body state or create appropriate motor actions or corrections. Note also that *α*-*γ* co-activation requires signals to arrive at the same time to the *α*- and *γ*-MN pools of a same muscle via *different pathways with different conduction velocities* (i.e., predominantly cortico-and proprio-spinal for *α*-MNs vs. cerebello-, reticuo-, rubro-spinal tracts and brainstem vestibular outputs for *γ*-MNs [63–68]). Lastly, *α*-*γ* co-activation only biases the *presynaptic* input to the *α*- and *γ*-MN pools, but does not directly provide the *γ*-MNs with the *actual postsynaptic α*-drive to muscle fibers, as mentioned below.

### A humble homonymous *α*-to-*γ* collateral performs as well as idealized *α*-*γ* co-activation

The main contribution of this work is that it confronts us with the previously unknown true cost of unmodulated velocity-dependent stretch reflexes, while also proposing an alternative to *α*-*γ* co-activation that is evolutionary and physiologically plausible at the level of a homonymous *α*-to-*γ* collateral.

Such collateral projection among MNs have long been observed in studies of the cat and mouse spinal cord (Fig. 8A) [31–34, 47], but not interpreted in this context, or for this functional role. Rather, the functional role of that reported inter-motoneuronal facilitation was only speculated on and interpreted as connections among *α*-MNs. Importantly, those studies [33, 47, 69, 70] did not confirm or deny that the collaterals were from *α*-MNs to *γ*-MNs as we propose here. Thus, prior studies partly support our proposed mechanism, even if their experimental limitations could not conclusively identify projections to *γ*-MNs. However, we believe that it is not unreasonable to suppose that such functional collateral projections to *γ*-MNs indeed exist. In addition, recent computational work also argues that Ia afferent signals for voluntary movement require fusimotor modulation independent of corticospinal drive [30]. We believe our mechanism can provide such modulation. Experimental validation of our proposed circuit will require the maturation of some promising optogenetic techniques that could show such low-level control of *γ*-MN pools in behaving animals [34, 71, 72].

Our proposed spinal level mechanism for scaling *γ*_*dynamic*_ MN activation by the homonymous *α*-drive collateral is not only generalizable to any novel or learned movement, but also independent of the cortical, subcortical or propriospinal competition at the presynaptic *α*-MN level. Similar to some of the limitations of *α*-*γ* co-activation and its variants mentioned above, conduction delays, pre-synaptic competition and inhibition can affect the homonymous *α*-to-*γ* collateral. Understanding how such effects arise within the intricate spinal circuits will require further work.

The advantage of an homonymous *α*-to-*γ* collateral is that it projects the actual (i.e., *postsynaptic*) *α*-MN drive to muscle fibers. As such, this modulatory mechanism to the *γ*-MN sidesteps the uncertainty arising from the *presynaptic* synthesis and competition among cortical, subcortical or propriospinal presynaptic projections to *α*-MN pools that idealized *α*-*γ* co-activation must consider.

We were careful to make an explicit quantitative and statistical comparison between idealized *α*-*γ* co-activation and our proposed homonymous *α*-to-*γ* collateral (Fig. 3), as shown in Figures. 6 and *SI Appendix*, Fig. S2. These results show that both approaches have functionally equivalent, though not identical, performance. This supports the face validity of *α*-*γ* co-activation that has been a fundamental tenet of sensorimotor neuroscience but, as mentioned above, is of uncertain origin.

### Locally-mediated modulation of *γ*-MNs via *α*-MN collaterals enables meaningful cerebellar and cortical learning and adaptation mechanisms

Biological and machine learning have the fundamental requirement that the system in question be minimally controllable, observable and predictable [73, 74]. Said differently, *meaningful error signals* are necessary for any effective and efficient learning processes. Our results for unmodulated velocity-dependent stretch reflexes for voluntary movement show that a realistic multiarticular limb with afferented muscles will exhibit disruptions that are movement-specific, typically large and variable, and that could even change movement direction as the velocity-dependent reflex gain increases. Therefore, unmodulated velocity-dependent stretch reflexes present any learning strategy with error signals that are at best highly nonlinear, and at worst not meaningful for learning. This makes it difficult, or even impractical, to learn limb movements from a naïve state, or replicate learned movements that are not identical each time they are performed. Placing our results in the context of the rich literature on motor learning and control, and using cerebellar circuits as an example, we argue that the regulatory effects the proposed homonymous *α*-to-*γ* collateral *at the spinal level* in fact serve as a critical enabler for learning, performance and adaptation. This complements recent evidence that spinal sensorimotor adaptation and learning mechanisms can be enabled by Renshaw cells [75].

Current thinking is that computational frameworks of the cerebellum favor hierarchical reinforcement learning with predictions via multiple internal models [55]. However, forming, refining and exploiting an internal model of any variety from a naïve state requires experience with a minimally controllable, observable and predictable system. We propose that this low-level circuit for locally-mediated modulation of *γ*-MNs via *α*-MN collaterals regularizes any new voluntary limb movement to the point that it can *enable learning* from a naïve state by combining motor babbling [76] or directed practice [77] with a higher-level learning strategy. It is to the advantage of the individual to be born with a body that is minimally controllable from the start.

As can be seen from the cumulative and terminal errors in Figure 5C, this low-level circuit is far from a panacea for all movements—and leaves room, and need, for improvements via additional supra-spinal mechanisms. Note that sample movements that show large disruptions after reflex modulation still have typical muscle activation changes *SI Appendix*, Fig. S3. This collateral circuit could serve as a low-level regulator that enable exploration-exploitation during the formation of an internal model (or Bayesian priors, synergies, gradient-descent strategies, etc. if the reader is not of the internal-model persuasion [41]). From an evolutionary perspective, we could even speculate that such a low-level circuit is an ancient enabler of movement as primeval *β* skeleto-fusimotor MNs in amphibians and reptiles (e.g., a tricycle) led to separate and independent *α*- and *γ*-MNs in mammals [13] (e.g., a bicycle)—and the need arose for some initially effective and parsimonious form of *α*-*γ* coordination at the spinal level (e.g., training wheels on the bicycle) for the control of limb impedance before higher circuits evolved or are trained. We speculate that, like *β*-MNs, the proposed collaterals are the *afferentation Yin* that complements the *efferentation Yang* of Hennemann’s Size Principle to enable low-level, robust regulation of arbitrary movements. These collaterals could collaborate with the posited *α*-*γ* co-activation, and the few *β*-MNs in mammals, to create a flexible *fusimotor ecosystem* that enables voluntary movement. By locally and automatically regulating the highly nonlinear neuro-musculo-skeletal mechanics of the limb, this low-level fusimotor circuit collaborates with, and enables, high-level brainstem, cerebellar and cortical mechanisms for learning, adaptation, and performance. In fact, as ontogeny recapitulates phylogeny, *β*-MNs that are followed by homonymous *α*-to-*γ* collateral as early regulators of velocity-dependent stretch reflex during an individual’s development could in time be complemented by more sophisticated fusimotor controllers as they become available (such as those reported and intensely studied for cerebellar control of movement [55]).

### Limitations and future work

The scope of this computational study is limited to the investigation of the disruption of voluntary movement caused by velocity-dependent stretch reflex from Ia afferent nerve fibers. Our spindle model is an over-simplified version of previously described models [7, 78, 79]. Moreover, we assume that there is appropriate *γ*_*static*_ MN drive that keeps the muscle spindle from going slack. Thus we do not consider stretch reflex signals from II afferents, or tendon tension signals from Golgi tendon organs [37, 38]. Another key limitation is that our spinal circuit is simplified in the extreme and does not consider the divergent and convergent branching of Ia, Ib and II afferent signals to multiple homonymous and heteronymous interneuronal pathways within and across muscles and limbs, where monosynaptic inputs to even antagonist motor neuron pools are largely overlapping [37, 80, 81], and even the *α, β, γ* classification of MNs is evolving [82]. Future work is needed to further our investigations of the fusimotor system. Similarly, we use a simple Hill-type muscle model included in MuJoCo, which can be improved by recent work (e.g., [83]).

Lastly, we necessarily present the best-case scenario for the mitigation of cumulative and terminal errors as we do not consider mono- and di-synaptic time delays in our propose modulation of *γ*-MN activity. However, time delays are also an unexplored and unresolved issue in idealized *α*-*γ* co-activation and its variants. Future work can address conduction and computational delays, as well as nonlinearities and delays from recruitment and rate-coding, muscle activation-contraction dynamics [84], etc.

From a behavioral perspective, our simulated tasks are not meant represent a specific task-related upper limb movements such as reaching or joint flexion/extension [19, 30, 39, 46, 85]. Rather, we start with open-loop arm movements that explore the 3D workspace so as to ask the fundamental question of the effects of disruptions from velocity-dependent stretch reflex in general. Nevertheless, it is worth considering whether the effects of velocity-dependent stretch reflexes on the simulated movements can extend to movements of functional importance for humans, and especially reaching movements compromised by pathologic synergies in neurological conditions such as stroke, or tremor in Parkinson’s disease. For this, it will be necessary to incorporate more detailed models of the muscle spindle, spinal circuitry, and tasks relevant to human functions—and of the neuropathology of interest.

## Methods

### Movements without velocity-dependent stretch reflex feedback

We created 1,100 open-loop three-dimensional arm movements of a Rhesus Macaque (Macaca mulatta) arm model, each lasting two seconds with a 2000 sampling rate. The model was adapted from the SIMM (Musculographics Inc) model developed by Moran et al. [86] into a MuJoCo model (**Mu**lti-**Jo**int dynamics with **Co**ntact) by first loading the SIMM model into an OpenSim (Open Source Simulation and Modeling) model [87] and then converting the OpenSim model into MuJoCo [88]. The adapted MuJoCo model is shown in (Fig. 1C) with the same body segment lengths, joint limits, and tendon routing as the original model. The number of muscles and degrees-of-freedom (DOFs) were reduced, respectively, to 25 muscles and 5 DOFs (shoulder abduction/adduction, shoulder flexion/extension, shoulder rotation, elbow flexion/extension, and forelimb pronation/supination). We excluded hand muscles and fixed the wrist joint as they are unnecessary for the simulated arm movements. The musclotendon model in MuJoCo is a Hill-type with inelastic tendons [89]. The muscle force parameters and tendon slack lengths were set as in the original model.

Each of the 25 Hill-type muscles received a feed-forward *α*-MN drive signal (Fig. 1A), whose level could vary from zero to one, which maps to 0% to 100% muscle activation and muscle force [90]. The feedforward *α*-MN drives were created as a beta probability density function to generate beta shapes which then were scaled and transformed into ramp signals. In each arm movement, five randomly-selected muscles were activated from zero to 60% of maximum (Fig. 1B), while the remaining 20 muscles reached only 4% of maximum muscle activation (inset). This selection of *α*-drive signals assumed a random exploration that did not consider motor primitives, synergistic control or any optimization strategy (see Discussion). This distribution of high and low activations mitigated co-contraction and enabled both small and large arm movements with maximal endpoint displacements ranging 5.178 cm to 46.87 cm that spanned the full workspace of the 47.35 cm length arm model (*SI Appendix*, Fig. S1). The trajectory of the endpoint (distal head of the third metacarpal) of the open-loop arm movements served as reference endpoint trajectories (Fig. 1C) for computing deviations of the endpoint trajectory of arm movement with velocity-dependent stretch reflex feedback from the open-loop endpoint trajectories.

### Movements with unmodulated and modulated velocity-dependent stretch reflex feedback

Excitatory velocity-dependent (Ia afferent) stretch reflexes from muscle spindles form feedback loops to alpha-motoneurons of the homonymous extrafusal muscle via spinal pathways [37]. We added a simple muscle spindle model to each of the 25 Hill-type muscles of the macaque arm. The model takes muscles velocity input and generates Ia afferent as positive muscle velocity (i.e., velocity-dependent stretch reflex) output. For each of the 1,100 arm movements, we performed closed-loop simulations of the movement with the velocity-dependent stretch reflex feedback of different reflex gain *k* from 1 to 10. We show these gains are physiologically tenable by computing peak change in muscle activation caused by the velocity-dependent (Ia afferent) stretch reflex feedback (Fig. 7) and compare them to reflexes elicited in human arms (up to 40%MVC reflex EMG) during interactions with destabilizing environments [39].

The neural circuit, schematic diagram of the closed-loop simulation of arm movements with unmodulated velocity-dependent stretch reflex gains, are shown in figure (Fig. 1, bottom plots). The descending commands were the same as feedforward descending commands (DSC) to *α*-MN drive during the open-loop simulation. The muscle spindle of each muscle received the muscle velocity (*V m*) as input and generated the stretch velocity of the muscle (*V*_*stretch*_) as positive muscle velocity for when the muscle was lengthening or zero for when shortening or isometrically contracting (i.e., negative velocity). The muscle stretch velocity was then multiplied by an unmodulated reflex gain *k* to produce the velocity-dependent stretch reflex feedback (*k*\**V*_*stretch*_). The muscle activation (*α*-drive) was computed as follows:

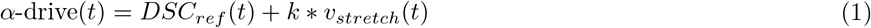

Where *DSC*_*ref*_ (*t*) refers the feedforward descending commands to *α*-MNs at time *t* and *v*_*stretch*_ is positive muscle velocity in lengthening muscles and zero in shortening or isometrically contracting muscles.

We investigated how modulation of velocity-dependent stretch reflex either by implementing an idealized *α*-*γ* co-activation (Fig. 3A) or via homonymous *α*-to-*γ* collateral (Fig. 3B) changes the disruptions in the endpoint trajectories. For the idealized *α*-*γ* co-activation simulation, the descending command to *α*-MN of the muscle scaled its *γ*-MN reflex gain *k*(Eq.2) while for homonymous *α*-to-*γ* collateral simulation, the output of the *α*-drive of the muscle scaled its *γ*-MN reflex gain *k* (Eq. 3). Multiplying the reflex gain *k* (an integer ranging from zero to ten) by the descending command or *α*-drive (a value between zero and 1) was similar to tuning down the tonic gain of *γ*-MN. The muscle activation for idealized *α*-*γ* co-activation was computed as follows:

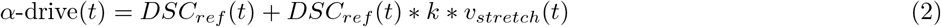

and for homonymous *α*-to-*γ* collateral:

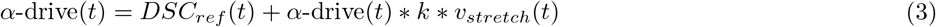

Similar to open-loop simulations, we recorded trajectories of the endpoint at each gain *k* and computed deviation in the movement trajectory (i.e., cumulative residual, CR) and deviation in of terminal position (i.e., terminal error, TE) of the endpoint trajectories from their reference endpoint trajectory of the open-loop arm movement. CR is the mean of the Euclidean deviations in the movement trajectory(Eq. 4):

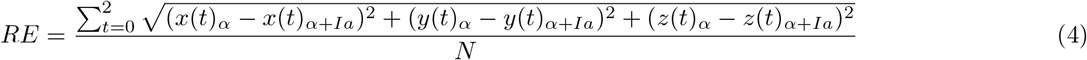

and TE is the deviation of the terminal position of the endpoint(Eq. 5):

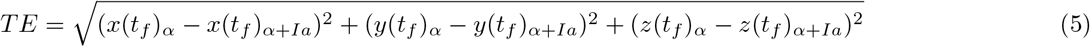

The x,y,z positions of the endpoint for open-loop arm movements(i.e., movements without feedback) are *x*(*t*_*f*_)_*α*_, *y*(*t*_*f*_)_*α*_, *z*(*t*_*f*_)_*α*_ and *x*(*t*_*f*_)_*α*+*Ia*_, *y*(*t*_*f*_)_*α*+*Ia*_, *z*(*t*_*f*_)_*α*+*Ia*_ for movements with velocity-dependent stretch reflex (Ia afferent) feedback at a reflex gain *k. N* is the total number of samples (two seconds at 2000 sampling rate). The magnitude of the disruption of the arm endpoint trajectory at each gain was quantified by scaling CR and TE of each movement to its to maximal endpoint displacement (*SI Appendix*, Fig. S1). We quantitatively compared the cumulative distance between the endpoint trajectories of arm movements with reflex gain modulated as per idealized *α*-*γ* co-activation (Fig.5B) and scaled via the homonymous *α*-drive (Fig.5C) using dynamic time warping (an inbuilt MATLAB function “dtw”). All analysis and statistical procedures were performed in MATLAB using non-parametric statistics Wilcoxon-Mann-Whitney. A Bonferroni correction was applied to adjust for multiple comparisons.

## Acknowledgments

The authors would like to thank Junhyuck Woo for his assistance with initial development of the computational model in MuJoCo in the course BME 504, and Drs. Kazuhiko Seki, Gerald Loeb, Sten Grillner, and John Krakauer for their insightful comments on earlier versions of this work. We thank the anonymous reviewers for their critical and useful comments.

## Supporting Information Appendix

**SI Appendix, Fig. S1.**
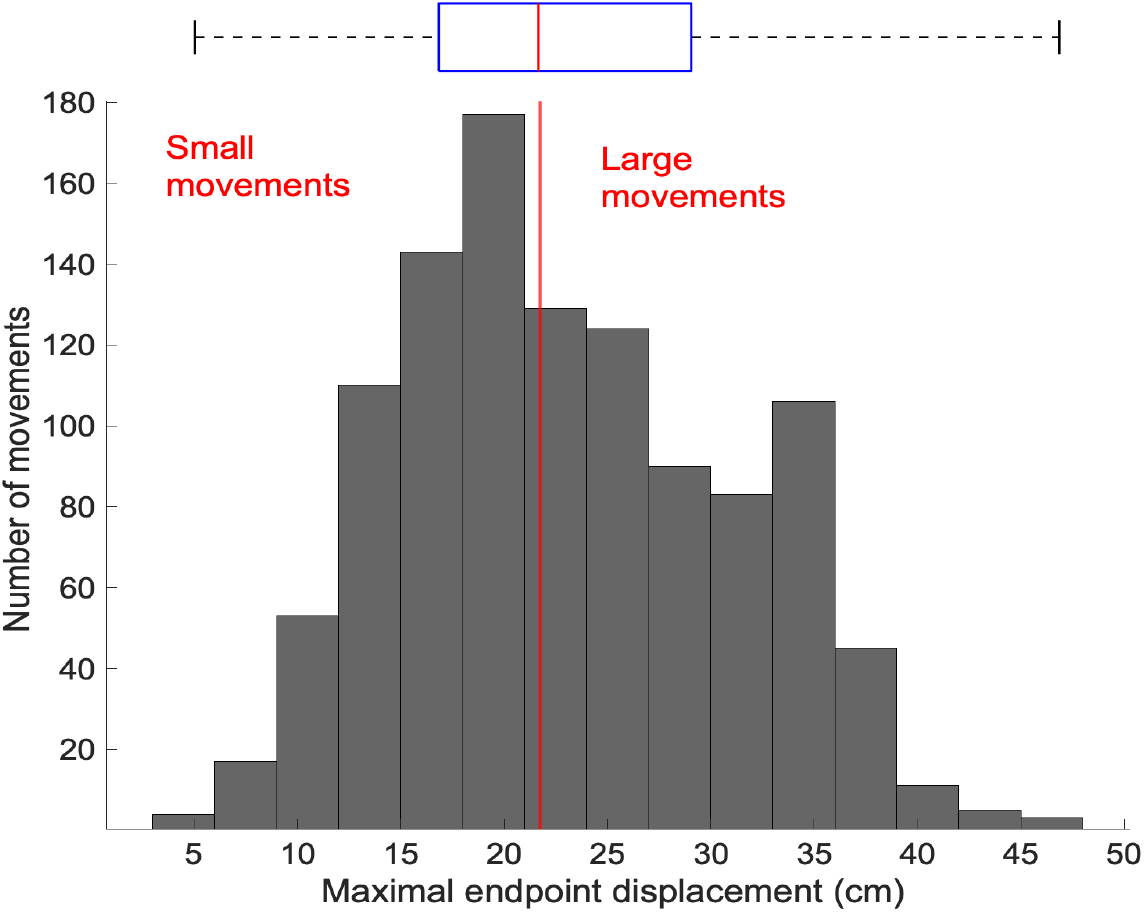
Arm movement amplitudes. Distribution of maximal endpoint displacement of the reference trajectories of the open-loop movements.

**SI Appendix, Fig. S2.**
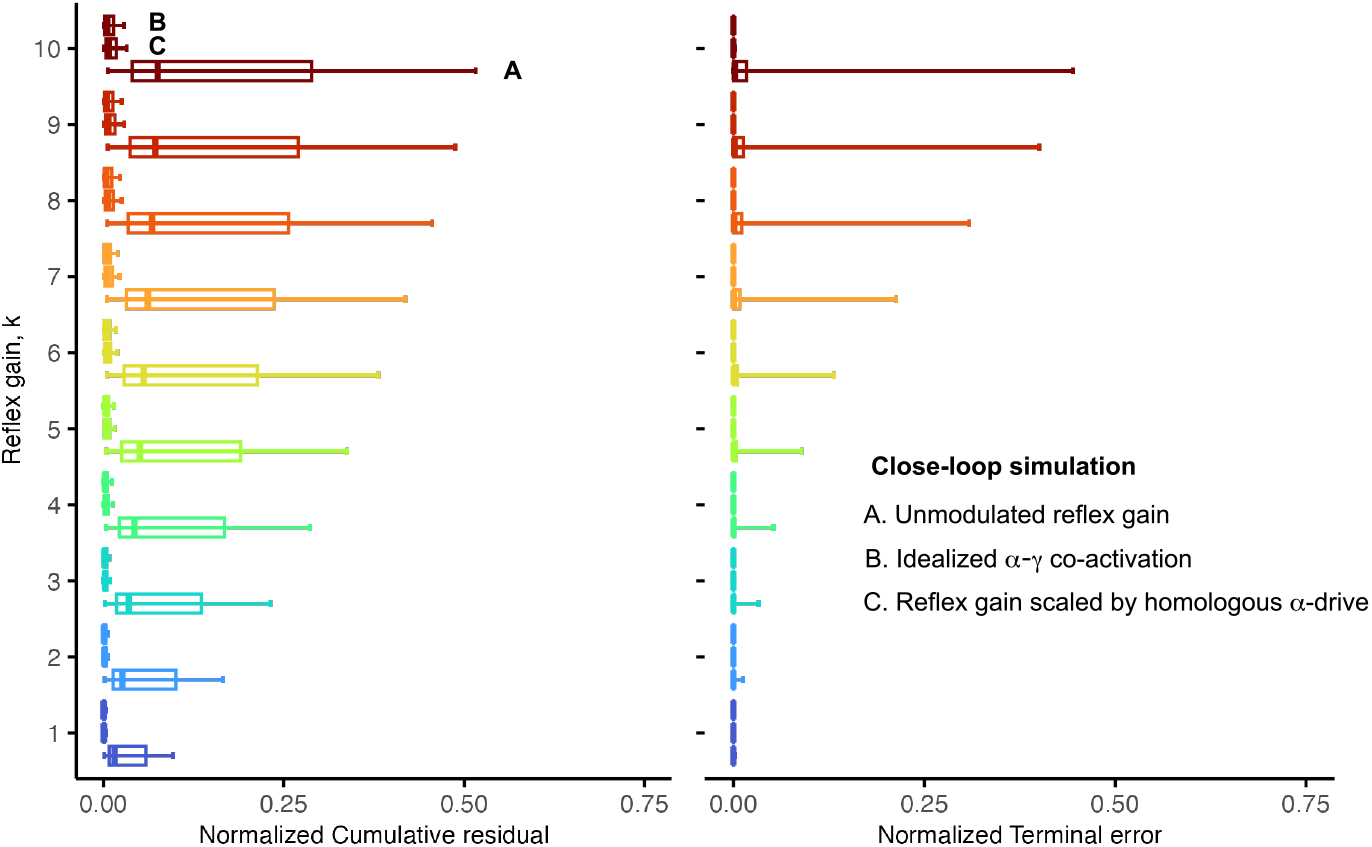
Distribution of cumulative residual and terminal errors shown in Figure 5. The rightmost whisker shows the 90th percentile of the data.

**SI Appendix, Fig. S3.**
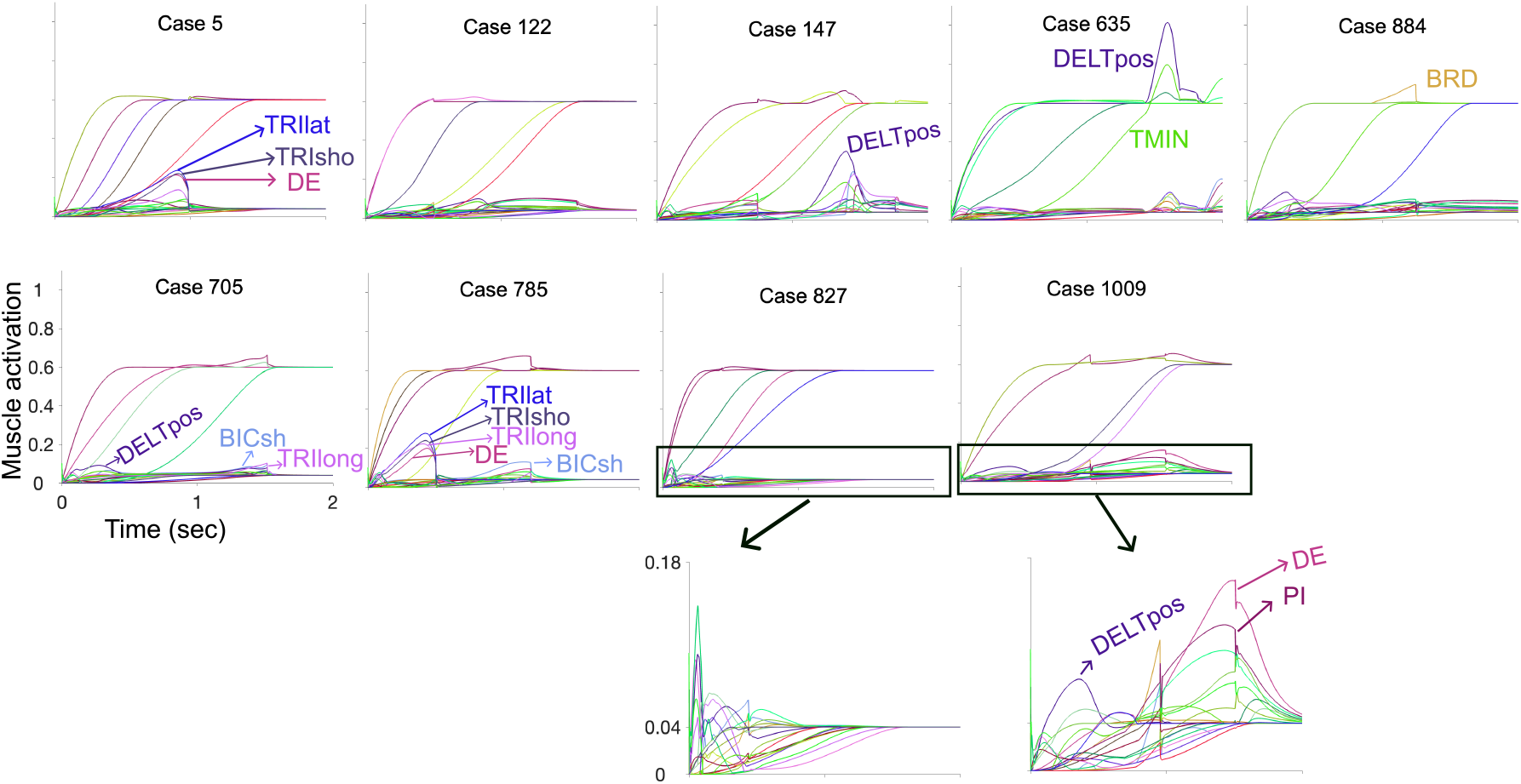
Sample muscle activation with velocity-dependent stretch reflex feedback at maximal reflex gain. Nine examples of *α*-MN-drive to muscles during closed-loop simulation with velocity-dependent stretch reflex at a reflex gain *k* of 10. Top plots are examples of cases shown in fig.2 in which scaling the velocity-dependent stretch reflex by the *α*-MN-drive to each muscle significantly reduced disruption in the movement trajectory and terminal position. Bottom plots are examples of cases shown in fig. 4 that had large disruptions even when the velocity-dependent stretch reflexes are scaled by the *α*-MN-drive to each muscle. Some representative muscle labels are shown, as defined in [86].

